# Revised Adaptive Immune Receptor Data in the Immune Epitope Database

**DOI:** 10.64898/2026.06.03.728549

**Authors:** Lonneke Scheffer, Eve M. Richardson, Randi Vita, Laura Zarebski, Nina Blazeska, Daniel K. Wheeler, Jason R. Cantrell, Sebastian N. Deleuran, William D. Lees, Scott Christley, Brian Corrie, Lindsay G. Cowell, Alessandro Sette, Bjoern Peters

## Abstract

The Immune Epitope Database (IEDB, iedb.org) is a freely available resource that catalogs experimentally defined immune epitopes and – if available – the immune receptors that recognize them. Currently, the IEDB records ∼185, 000 T cell receptors and ∼5, 000 B cell receptors/antibodies with experimentally verified epitope specificity. Because these receptor data were manually curated from ∼3, 300 references spanning decades, nomenclature inconsistencies present challenges for computational analyses and user queries. To support integrated analysis of the entire dataset, we revised the IEDB receptor data standardization and validation pipeline to flag and correct inaccuracies. Anomalous receptors from over 800 studies were flagged for re-curation. The updated receptor dataset shows greater conformity through consistent gene nomenclature formatting and harmonized CDR sequence delimitation. Taking advantage of the increased receptor data consistency, the IEDB web interface was expanded to include receptor search features directly on the homepage, support V/J gene and species options in the refined receptor search, and allow direct data export in the Adaptive Immune Receptor Repertoire (AIRR) format. We anticipate that the improved receptor data quality will simplify bioinformatics analyses, and facilitate integration of IEDB data into cross-repository data resources, such as the AIRR Knowledge Commons.

## Introduction

T cell receptors (TCRs) and B cell receptors (BCRs / antibodies) form a highly diverse class of molecules referred to as adaptive immune receptors, which enable the immune system to distinguish self from non-self antigens and initiate responses to potential threats. The diversity of immune receptors is driven by the stochastic V(D)J recombination process during which one of several variants of variable (V), diversity (D), and joining (J) gene segments are combined with a constant (C) region to form a receptor chain gene (1). Two chains assemble to form the immune receptor heterodimers, with BCRs consisting of heavy and light chains, and TCRs containing α and β or γ and δ chains. Each recombinant chain includes a set of three Complementarity Determining Regions (CDRs) which determine the receptor specificity, with the CDR3 being the most variable (1). Advances in next-generation sequencing of Adaptive Immune Receptor Repertoires (AIRRs), as well as immunological assays for functional validation of immune receptors have resulted in increasingly large datasets rapidly becoming available (2–4).

For over two decades, the Immune Epitope Database (IEDB, iedb.org) has curated and catalogued experimentally determined epitopes in the context of infectious disease, allergy, autoimmunity, and transplantation (5), while the sibling database CEDAR (cedar.iedb.org) specializes on epitopes in the context of cancer (6). If experimentally determined, the IEDB also curates the sequence and 3D structure of the receptor recognizing the epitope (7, 8). This curated catalog of epitope-specific immune receptors has proven invaluable for the development and benchmarking of receptor specificity prediction tools (9–25). In 2023, the IEDB joined the AIRR Knowledge Commons (AKC) project (26), which aims to integrate repositories holding complementary immune receptor repertoire, specificity, and germline gene data. The integration encompasses: (i) large-scale repertoire data from the AIRR Data Commons (27) through VDJServer (28) and iReceptor (29), (ii) receptor-epitope specificity data from the IEDB and antibody-antigen specificity data from the Immune Receptor Antigen Database (IRAD, irad.roskinlab.org), and (iii) germline V/D/J gene information from the Open Germline Receptor Database (OGRDB) and accompanying population-level genotypes and haplotypes from VDJbase (30). The AKC will facilitate streamlined querying of data across multiple repositories, for instance by annotating receptors in large-scale repertoire datasets with experimentally validated or inferred epitope specificity. The foundation of this integration is the use of harmonized data representations and shared nomenclature across repositories.

Immune receptor sequence annotations are stored in the IEDB in two forms: *curated* and *calculated* data fields, both of which are available for export. The *curated* are human-curated, and intend to capture values as they were reported in the original publication (31). For TCR data, typically only the CDR3 of either one or both chains is reported, in addition to V and J gene names, while for antibodies, full paired V(D)J rearrangement amino acid sequences are often available. The format of the *curated* data is highly heterogenous due to the fact that it was derived from over 3, 000 articles by diverse authors spanning several decades during which V/D/J gene naming conventions have changed (32, 33), and CDRs have been reported according to differing amino acid numbering schemes (34–36). In 2018, computational procedures were introduced to compute gene usage, V domains, and CDR sequences from the *curated* V(D)J sequence data, which are stored as *calculated* data fields (7). However, these algorithms did not calculate data for all receptors, requiring users to combine *calculated* with the less-standardized *curated* data to construct the complete set. Further challenges arose from the inconsistent use of the term ‘CDR3’, which is often used colloquially to describe the sequence including the conserved Cys104 and Phe/Trp118 residues. However, in the IMGT definition these residues are excluded from the CDR3, and the segment that includes them is referred to as the IMGT Junction. The IEDB *calculated* CDR3 sequences followed the IMGT CDR3 format, whereas *curated* CDR3s were generally (but not always) reported as Junctions. This inconsistency led to users omitting CDR3s without the leading ‘C’ from their analyses (37), or attempting to correct data by prefixing/appending amino acids while relying on the error-prone assumption that exactly one residue may be missing on either side of the CDR3 (12, 14, 18, 38–40).

To improve the consistency, quality and usability of the receptor data in the IEDB, we revised our standardization and validation pipeline, and applied it to the full receptor dataset. A complementary set of computational tools were used to produce standardized, analysis-ready *calculated* data for all receptor records, such that they can directly be used in bioinformatics analyses without further reformatting. This consists of V, D and J gene names, as well as CDR1, 2, 3, Junction and V domain sequences. Specific attention was given to the standardization and validation of CDR3/Junction sequences, providing consistent coverage up to the anchor residues and concordance with annotated germline V and J genes. Furthermore, the improved data validation led us to detect and re-curate thousands of erroneous receptor records from over eight hundred studies, this quality control is now integrated into the curation process to ensure consistency of all future curation. Finally, we present enhanced receptor search functionalities in the IEDB web interface and added support for AIRR-compliant data export (41). We anticipate these changes will facilitate tool development and bioinformatics analyses, including epitope binding and immune response prediction studies, and further integration with the AKC.

## Overview and Scope of Immune Receptor Data in the IEDB

### Receptor Content of the IEDB

The IEDB organizes receptor data into ‘receptor groups’ by shared CDR3 sequences, receptor type (e.g., αβ TCR), and species, with the aim to group together similar receptors that are presumed to share the same specificity (7). As of to date, the IEDB contains 213, 100 epitope-specific B- and T cell receptors, which have been organized into 189, 386 receptor groups associated with 7, 831 epitopes. This corresponds to a tenfold increase in receptor data since August 2018 (7), with a notable surge in available T cell receptor data during 2020 as a result of the inclusion of a large-scale SARS-CoV-2 study (42) (see **Figure 1**). Nearly all of the TCR data in the IEDB is of human origin, with less than 4% of receptors originating from other species (primarily mouse) (see **Supplementary Table 1**). The antibody data stored in the IEDB spans across more diverse species, with approximately 60% of the receptors being of human origin and 12 species contributing more than 10 receptors each (see **Supplementary Table 2**).

**Figure 1:**
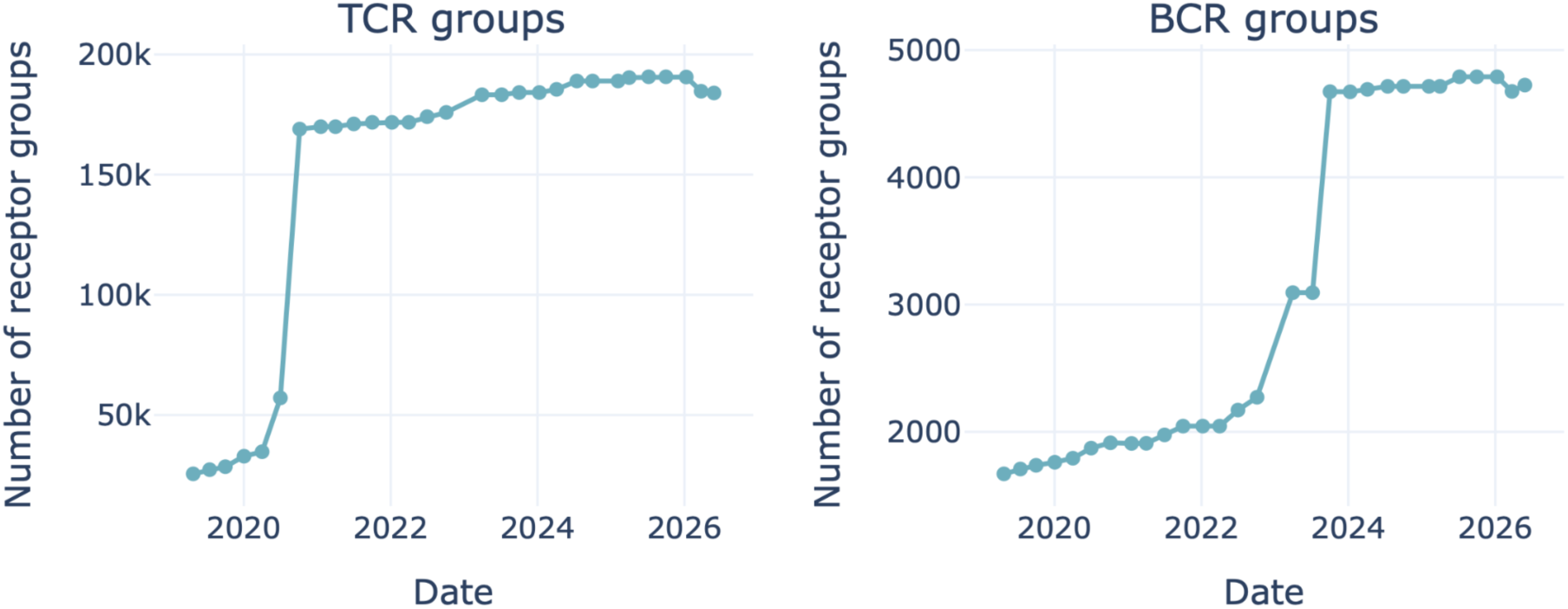
The increase in T and B cell receptor groups in the IEDB since 2019. The slight dip in receptor groups early 2026 is due to the merging of receptor groups with identical CDR3 sequences as a result of standardization efforts described in this work.

### Comparison to Other TCR Specificity Databases

Other frequently cited databases of curated TCR specificities are VDJdb (43) and McPAS-TCR (44). Between these databases, the IEDB provides the largest set of TCRs with known specificity described in the literature, due to curating both high- and low-throughput data (see **Table 1**). Furthermore, the IEDB integrates the receptor data into an extensively annotated epitope and assay context. This makes it possible to e.g., query for antibodies that are likely to be neutralizing (select B cell assays with ‘biological activity, neutralization’), or find out what cytokines are reported to be produced by T cells with a given epitope target (in the Assay export column: ‘Assay, Response measured’). While the IEDB and VDJdb are actively maintained as of spring 2026 (github.com/antigenomics/vdjdb-db, accessed 05/28/2026) and both provide programmatic access via public APIs, McPAS-TCR has not been updated since September 2022 (friedmanlab.weizmann.ac.il/McPAS-TCR, accessed 05/28/2026) and does not provide an API.

**Table 1:** Between the IEDB, VDJdb and McPAS-TCR, the IEDB stores the largest set of epitope-specific TCRs. Unique TCRs are defined based on their ‘receptor group’ criteria (CDR3 sequences, species and receptor type). Only αβ TCRs with peptidic epitopes are included in this comparison, which is the majority of the receptor data in the IEDB (see **Supplementary Tables 1 and 2** for the total numbers of receptors). The number of associated PubMed IDs (PMIDs) is shown, but TCR-epitope pairs without PMIDs are not omitted from the other columns as VDJdb and McPAS-TCR also include receptor data without PMIDs. Comparisons were performed using the latest data releases at the time of writing (IEDB: 05/28/2026, VDJdb: 05/16/2026 and McPAS-TCR: 09/10/2022).

| Database | Unique $\alpha\beta$ TCR-epitope pairs | $\alpha\beta$ TCRs with cognate peptidic epitope | Peptidic Epitopes with cognate $\alpha\beta$ TCR | Associated reference PMIDs |
| --- | --- | --- | --- | --- |
| IEDB | 195,808 | 181,168 | 2,849 | 763 |
| VDJdb | 115,459 | 111,941 | 1,792 | 337 |
| IEDB:VDJdb overlap (% of VDJdb) | 39,112 (33.9%) | 38,391 (34.3%) | 1,056 (59.2%) | 222 (65.9%) |
| McPAS-TCR | 14,433 | 14,023 | 370 | 116 |
| IEDB:McPAS-TCR overlap (% of McPAS-TCR) | 9,187 (63.7%) | 9,174 (65.4%) | 309 (83.5%) | 72 (62.1%) |

The fraction of receptors stored in VDJdb or McPAS-TCR which do not occur in the IEDB can generally be explained by the following scope restrictions: (i) the exact *epitope* is not known, and specificity is defined on the broader antigen or organism level, or (ii) the epitope is known but does not satisfy the inclusion criteria defined by the IEDB, such as epitopes failing to satisfy length/mass restrictions or HIV/SIV epitopes (IEDB Inclusion Criteria, curationwiki.iedb.org/wiki/index.php/IEDB_Inclusion_Criteria, accessed 05/28/2026), (iii) the data is not available as part of a published study. Aside from these defined restrictions, the IEDB team aims to curate all epitopes and cognate immune receptors described in the published literature.

## Receptor Data Updates

### An Improved Standardization Pipeline for Immune Receptor Data

The previous ‘2018 pipeline’ (7) employed ANARCI v1.1 (45) for V and J gene alignment and detection of the CDRs, and custom scripts additional processing. However, ANARCI does not support updating of the internally used V/J gene reference sets and requires amino acid sequence input, which allows less precise allele annotation than nucleotide sequence input. For the majority of TCR sequences in the IEDB, only CDR3 sequences and V/J gene names are reported (see **Supplementary Table 3**), necessitating additional standardization and validation steps for CDRs and gene names in the absence of the full sequence context.

An updated computational pipeline was implemented to ensure V domain and CDR subsequences are annotated according to the IMGT numbering scheme (35), and V/D/J genes are represented through IUIS/IMGT compliant names (32) (see **Figure 2**). For full sequence data, the pipeline employs IgBLAST (46) v1.22.0 for V/D/J gene annotation, which is run in ‘nucleotide’ sequence mode if available, otherwise in ‘amino acid’ sequence mode. For human BCR data, we used the human AIRR-C IG reference set provided by OGRDB (47) (IGH v9, IGK v4, IGL v3). Alleles with temporary names (i.e., those characterized by *i or substitutions such as *01_c330g) were excluded (6 IGHV and 5 IGLV alleles), thus only alleles with IUIS-approved names were retained. For human TCR data as well as mouse and macaque TCR and BCR data, we used the IMGT reference sets as included in the Immcantation suite v4.6.0 (hub.docker.com/layers/immcantation/suite/4.6.0). To detect V domains and CDR1, 2, 3 and Junction sequences from full sequence data we used ANARCII (48) v2.0.6 in ‘accuracy’ mode. As opposed to the former ANARCI algorithm used in the 2018 pipeline, ANARCII is species-agnostic and explicitly supports annotation of unconventional BCR sequences (shark VNAR or camelid VHH), allowing more sequences to be annotated.

**Figure 2:**
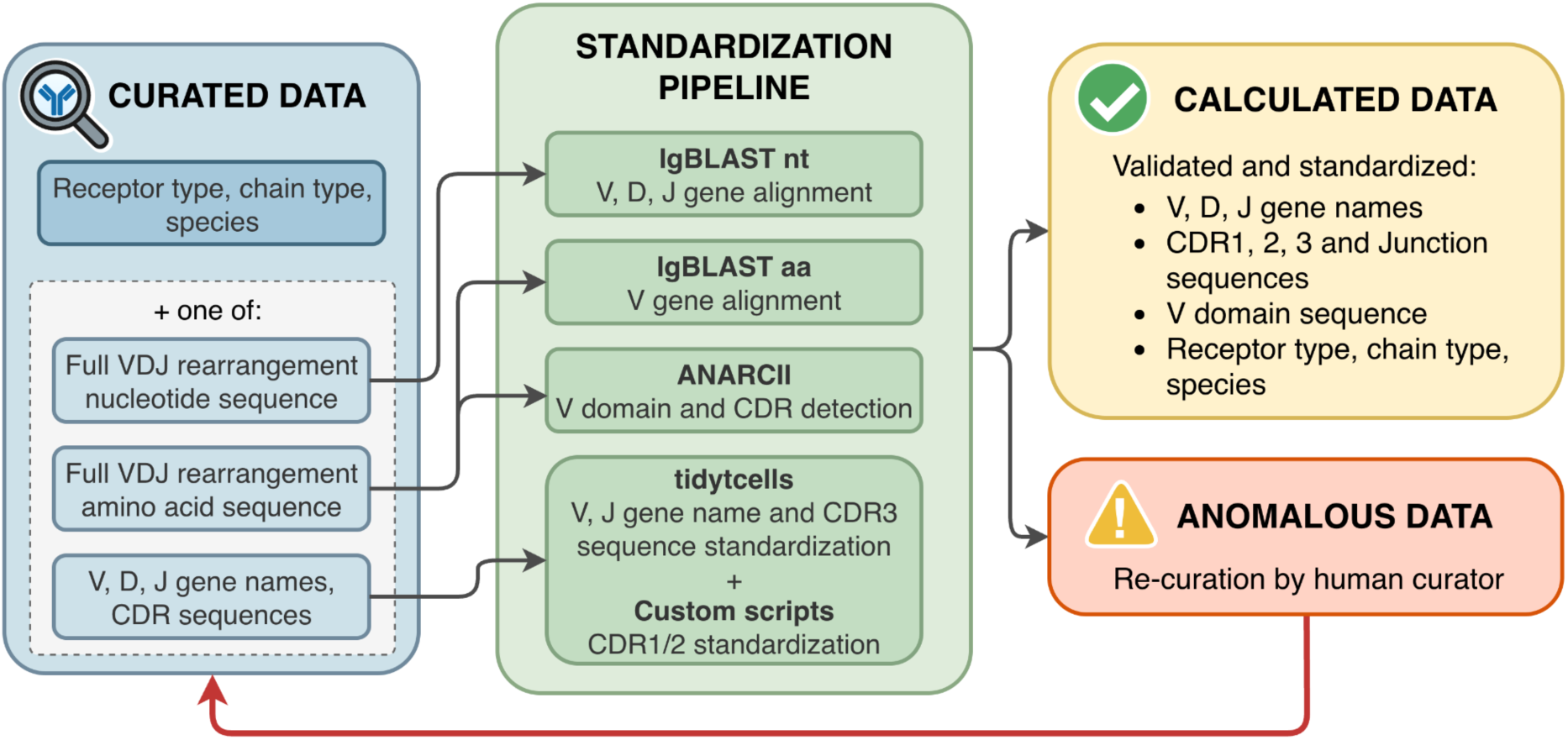
The updated receptor data standardization and validation pipeline. Depending on the format of the *curated* data (see **Supplementary Table 3**), a variety of computational tools are used to compute the *calculated* data output fields. Any failed tool runs or validation issues trigger the data to be flagged for re-curation.

For records where no full sequence information is available, we used tidytcells (39) v3.0.0-alpha (github.com/yutanagano/tidytcells/tree/dev/v3) to validate and standardize V and J gene names, and standardize CDR3 Junction sequences by comparison to the germline sequences of annotated V and J genes. *Curated* CDR1 and CDR2 sequences were validated by comparing to the IMGT germline CDRs of the *curated* V gene, if available. If the *curated* CDR was a subsequence of the germline CDR or vice versa, the sequence difference was presumed to be due to different numbering schemes (not true CDR sequence differences), and the germline CDR was adopted as the standardized sequence. When the *curated* and germline CDR contradicted (expected in BCR due to somatic hypermutations), no correction was applied. For TCRs only, the germline CDR1 and CDR2 sequences were included in the *calculated* data even if they were not curated, provided that sufficiently precise V gene information was available to unambiguously determine the CDR.

### Detection and Re-Curation of Anomalous Receptor Data

Extensive validation tests were implemented to detect and correct anomalous data in the IEDB, as well as to prevent such erroneous records from being entered in the future. During validation, we distinguish between *critical errors* and *warnings* based on severity and required intervention. Critical errors point out nonsensical *curated* data, and a manual correction of the curated record is required before any *calculated* data can be computed. Warnings are issued generously as consistency checks to detect potential anomalies and should prompt human review, but do not halt the computational pipeline and may be ignored on a case-by-case basis.

Critical errors are triggered by unresolvable species, unrecognized receptor and chain types, or invalid combinations thereof. Invalid characters in any of the input fields containing nucleotide or amino acid sequences result in a critical error as well. Unknown ‘X’ amino acids are permitted in the full amino acid sequences, but not in any of the CDR sequences. Warnings are raised when *curated* sequence data (including CDRs) are longer or shorter than expected, when there are mistmatches in the sequence data (e.g., a CDR not appearing in the full sequence), or when accession-derived sequences do not match curated sequences. Receptors were flagged for inspection when any of the computational tools raised warnings or failed to compute expected outputs, including low alignment e-values, invalid V/D/J gene names for the given species, CDR3 sequences that contradict the known reference sequence of the provided V or J genes (see **Figure 2**), or CDR1/CDR2 sequences mismatching their V gene germline sequence counterparts. *Calculated* genes and inferred loci are compared to *curated* chain types, which may reveal switched chain data or misidentified receptor types. Our data processing and tests are informed by the recommendations for developers using TCR germline references in (49).

The IEDB maintains and extensively utilizes ontologies and controlled vocabularies for consistent, accurate, and interoperable data representations (50). For human and mouse data, *calculated* V/D/J gene names are strictly limited to known IUIS/IMGT gene names at the allele, gene, or subgroup level, and consistent with the chain type and species. While host organism strain information is curated when available (Curation Manual2.0, curationwiki.iedb.org/wiki/index.php/Curation_Manual2.0, accessed 03/31/2026), the receptor chain species is set to the species which reflects the germline gene namespace. This means that for some receptors from TCR/BCR transgenic mouse strains, the receptor chain species is *Homo sapiens* to reflect the use of human V and J genes, and the transgenic mouse strain is stored as part of the assay information.

### Updates to the Curated and Calculated Receptor Data Fields

Running the validation tests on the total IEDB *curated* receptor data resulted in warnings or errors for over 14, 000 receptor chains. After human review of these issues, a total of 3, 008 receptor records (∼1.5% of the total receptor data) originating from 814 different references were marked for re-curation by revisiting the original publications and correcting the *curated* data fields. Subsequent to any corrections to the *curated* data, the updated standardization pipeline was used to (re)compute the *calculated* data fields for V/D/J genes, V domains, CDR1, 2, 3 and Junction sequences for all receptor data. For any field, the original *curated* data remain available, but *calculated* data, representing standardized values, is only provided if it could be unambiguously resolved and validated, ensuring high-quality data.

Most *calculated* TCR CDR3 Junction sequences remained unchanged from their *curated* CDR3 counterpart (84.9%, **Figure 3a**), whereas 9.5% required recovery of the conserved flanking residues, and 3.4% another type of correction. Non-standard or asymmetrical corrections typically involved adding the missing conserved Cys104 from the V gene, or removing an excess Gly119 from the J gene side (see **Supplementary Figure 1**). Most BCR CDR3 Junction sequences were recalculated from the full sequence, resulting in inclusion of the previously missing start and end amino acids (79.8%, **Figure 3a**). Because IEDB receptor groups are defined based on CDR3 sequences, these corrections required reassignment and consolidation of receptor group identifiers. As a result, the total number of receptor groups decreased by 6, 766 (see **Figure 1**).

**Figure 3:**
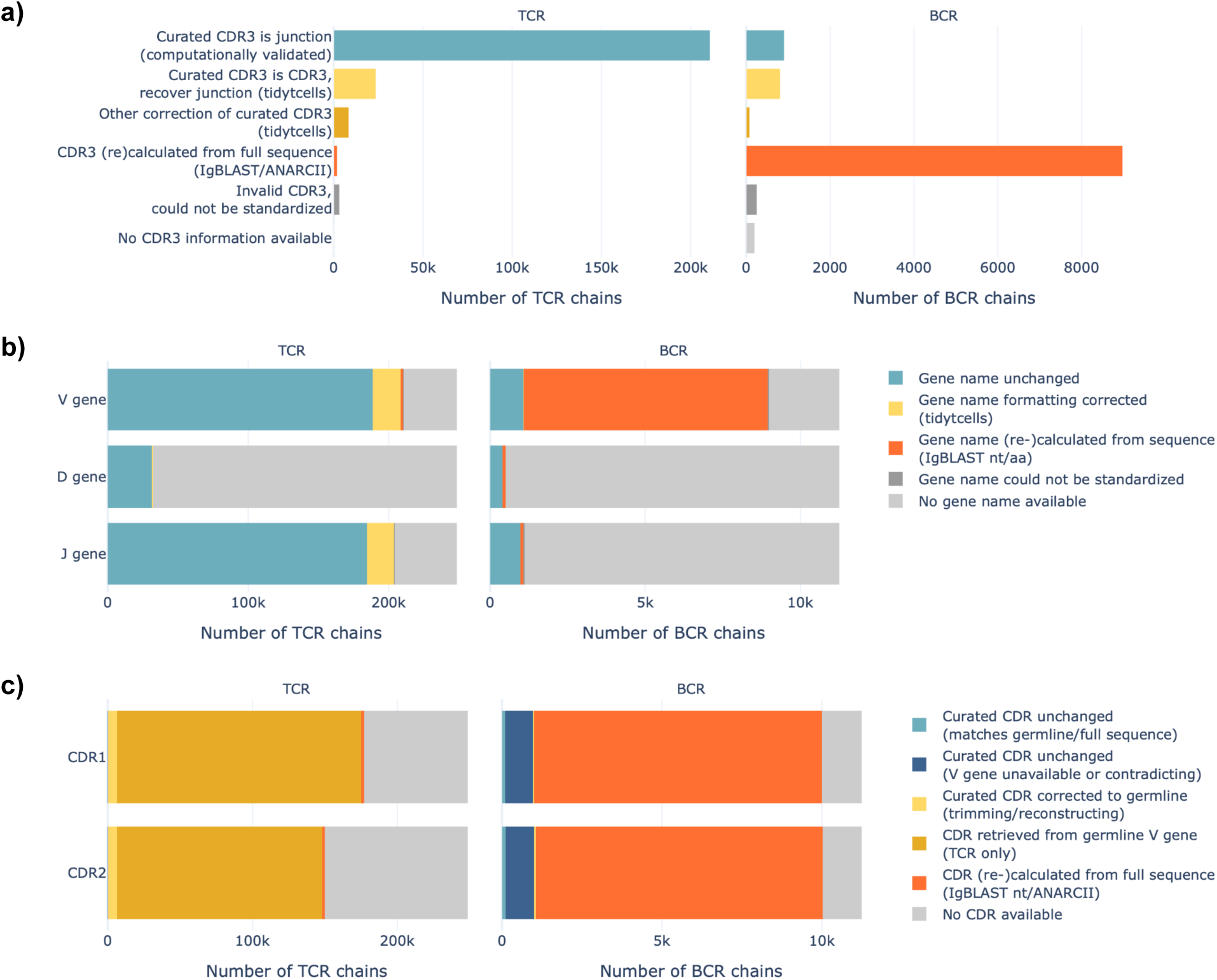
The standardizations applied to *curated* data to compute the *calculated* CDR and V/D/J gene data. a) The majority of *curated* TCR CDR3s corresponded to valid CDR3 Junction sequences, and a smaller fraction matched the IMGT CDR3 definition (no anchor residues), and were corrected. Other corrections of *curated* CDR3s (dark yellow) are described in more detail in **Supplementary Figure 1**. For most BCRs the CDR3 sequence could be computed from the full VDJ rearrangement sequence. Invalid CDR3s could not be standardized even after re-curation, and are kept only as *curated* CDR3 data. b) Some fraction of *curated* TCR gene names had non-standard formatting, such as ‘TCRBV01-01*01’ (corrected: ‘TRBV1-1*01’). All final names in the *calculated* data correspond to valid alleles, genes or subgroups for the given species. c) *Curated* V gene information was used to verify the *curated* CDR1 and CDR2 sequences. For most TCR data, CDR1 and CDR2 sequences were not directly curated but retrieved from the germline sequence of the *curated* V gene. Most of the BCR CDR1 and CDR2 sequences could be computed from full sequence information.

The *curated* TCR V and J gene data followed correct formatting in 75.9% and 74.3% of the TCR chains, respectively, and an additional 8.1% and 7.9% of the TCR V and J gene names could be standardized, e.g., correcting ‘TCRBV01-01*01’ to ‘TRBV1-1*01’. For BCR data, the V gene names were mostly (70.5%) recalculated from the full sequence (see **Figure 3b**). CDR1 and 2 sequences were mostly unavailable in *curated* TCR data, but recovered based on V gene information and added to the *calculated* data of 64.8% (CDR1) and 56.5% (CDR2) of receptor chains, respectively, and recalculated from full sequences for most BCRs (see **Figure 3c**). For BCR ‘light’ chains, we introduced the sub-types ‘kappa_light’ and ‘lambda_light’. These more precise light chain types could be inferred from IgBLAST V gene annotations or *curated* gene names for 97.1% of the BCR light chains.

### Improved Conformity of the Calculated Receptor Data

To assess how the re-curation and standardization efforts impacted the final *calculated* fields of the IEDB receptor dataset, we computed several overall metrics. The receptor data show greater conformity, as exemplified by a reduced number of V/D/J gene name representations, in particular for TCR data (see **Figure 4a**). Furthermore, the number of different CDR1, 2 and 3 sequences observed across chains with the same *calculated* V gene annotation was nearly always reduced in the standardized dataset (see **Figure 4b**). While for CDR3 sequences this effect can partially be explained by the omission of invalid CDR3s in standardized data, this is only a minor fraction (1.3% and 2.4% of respective TCR and BCR CDR3s, see **Figure 3a**). For CDR1 and CDR2, sequences were only standardized, but not removed (see **Figure 3c**).

**Figure 4:**
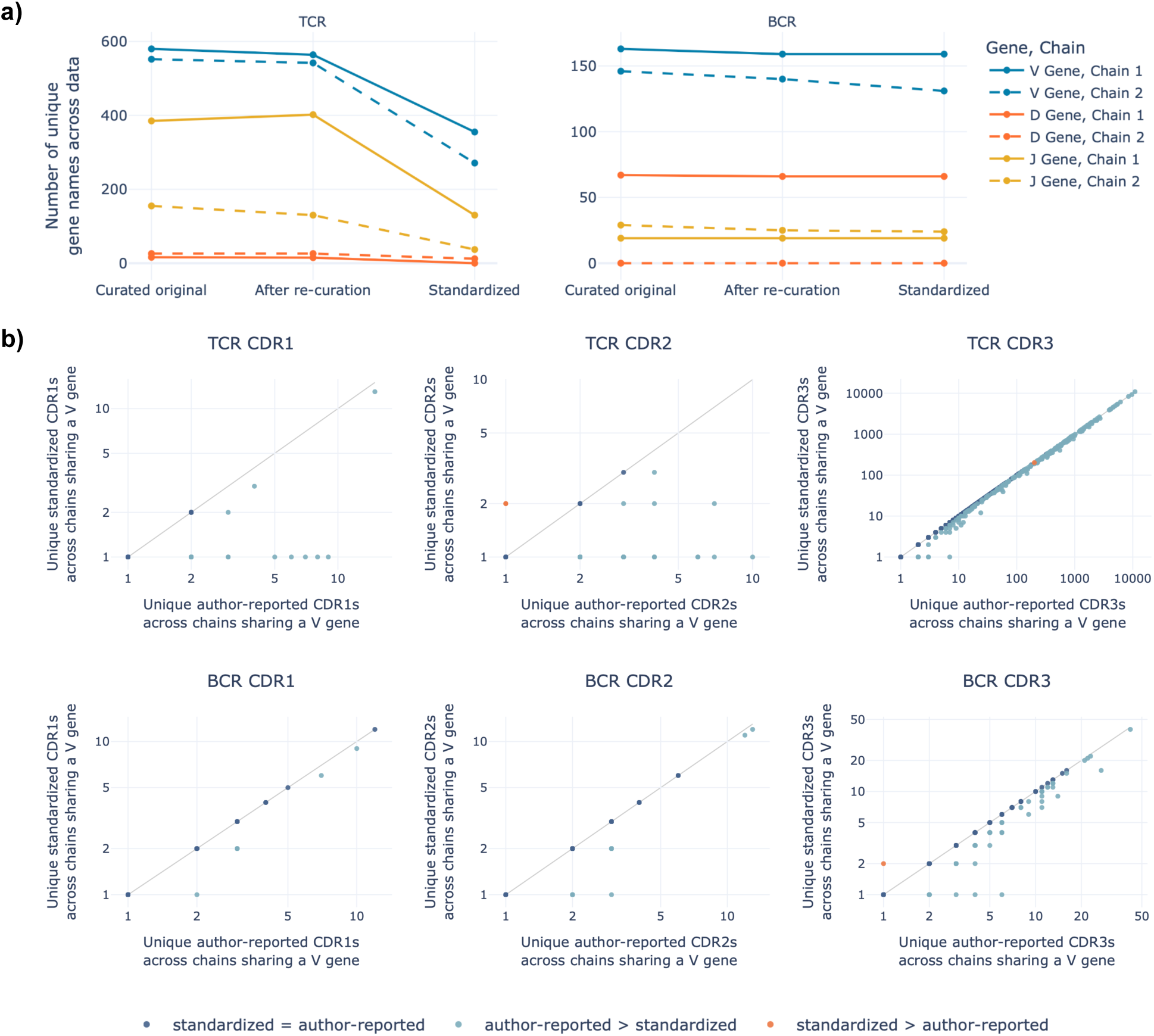
The standardized receptor data shows greater conformity. a) The total number of unique V/D/J gene names used across all receptors is reduced after standardization, in particular for TCR data. b) The total number of different CDR1, 2 or 3 sequences observed across all chains with the same *calculated* V genes was in nearly all cases reduced after standardization. The V genes in this plot represent each unique string value of the standardized gene names (at the gene, allele or subgroup level). For chains annotated with multiple V genes, CDRs are counted for each V gene. A more detailed comparison of TCR CDR3β sequences is given in **Supplementary Figure 2**.

As expected, the standardized CDR3 sequences are less diverse at the flanking regions due to better alignment (logoplot examples for TCR CDR3βs are shown in **Supplementary Figure 2)**. For the human antibody data re-annotated using the AIRR-C IG reference sets, the updated annotations were found to have significantly higher sequence identity to germline (see **Supplementary Figure 3**). This is particularly notable for kappa light chains, where 23.3% of annotations did not match a valid human IGKV gene, as genes from a mismatched species had originally been assigned.

### Enhanced Receptor Data Utility through Increased Matches

The standardization of the IEDB receptor data was a key prerequisite for integration into the AKC (26), which will simplify the matching of epitope-specific receptors from the IEDB to previously unannotated receptors from (disease-associated) repertoires. The AKC contains nearly ten thousand repertoires from the AIRR Data Commons, collectively containing over five billion unique receptor chain rearrangements. As of now, the αβ TCRs from the IEDB have been loaded into the AKC, yielding exact CDR3 matches across hundreds of TCR α and thousands of TCR β repertoires (see **Supplementary Figure 4**). Importantly, 5, 648 out of 31, 175 TCR groups with α-chain CDR3 matches and 10, 557 out of 100, 109 with β-chain CDR3 matches in the AKC had been subject to CDR3 standardization updates, meaning these matches were not possible prior to standardization.

As a complementary example of improved data matches for antibody data, we analyzed an external dataset from six subjects at baseline and 10 days post-RSV challenge from (51). We used CloneSearch (23) to predict Respiratory Syncytial Virus (RSV) specificity of antibody heavy chain sequences in six subjects at baseline and 10 days following RSV challenge, using the human RSV-specific antibodies from both original and updated IEDB datasets. Improved annotations of the IGHV gene name and CDRH3 sequence lead to better analytical outcomes, with the updated receptor dataset showing a higher proportion of sequences predicted to be RSV-specific at day 10. The median fraction of RSV-annotated antibodies increased from 0.05% in the original database to 0.33% in the updated database (p = 0.03, see **Supplementary Figure 5**).

## Web Interface Updates

### Receptor Search and Results Display

The ability to query the IEDB for receptor data and get consistent and complete results is greatly improved by the standardization effort. This encouraged us to increase the visibility and functionality of receptor queries (see **Figure 5a**). All interface changes described here have been mirrored in our companion site CEDAR (6). The Receptor (TCR/BCR) search box has been added to the homepage, allowing users to restrict their initial query to only include results where associated receptor sequences are available (‘has TCR/BCR’). This search box enables quick search for CDR3/Junction sequences or specific chain types. After the initial search query from the homepage, users can further refine parameters using the left-hand side panels (see **Figure 5b**). We added new settings to the refined receptor search panel for V/J gene usage and receptor species. Previously, the ‘Host’ species could be set, but this approach did not account for cases such as transgenic mice expressing human antibodies. With the new approach, the species can be restricted on the receptor level. Lastly, on the ‘Receptors’ results tab, a new ‘Epitope’ column is now displayed alongside receptor group information, making the receptor-epitope pairings clearer (see **Figure 5c**). It should be noted that any receptor search parameters can be freely combined with other filter options, allowing users to search, for instance, for a restricted receptor type (human TCR αβ with TRBV6-5) in combination with a certain epitope property (peptidic epitopes of length 10) presented on MHC Class I, all in one query. More detailed documentation of receptor data in the IEDB web interface can be found here: discuss.iedb.org/t/tcr-and-bcr-antibody-sequence-data.

**Figure 5:**
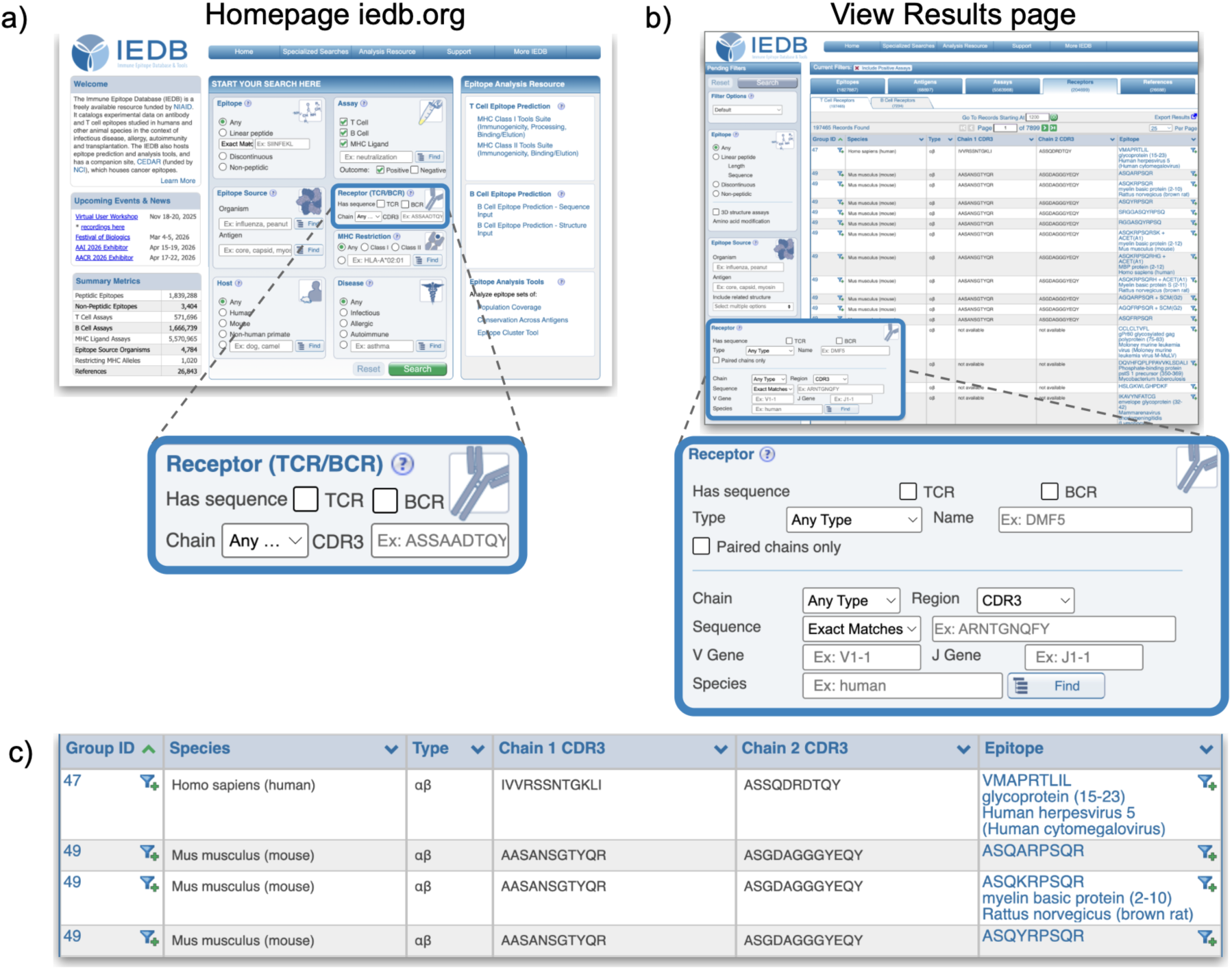
The updated receptor search and display functionalities on the IEDB web interface. a) The Receptor (TCR/BCR) search box was added above the MHC Restriction box on the homepage. b) The refined ‘Receptor’ search box on the results page now supports ‘V gene’, ‘J gene’ and ‘Species’ search. c) A new ‘Epitope’ column was added to the receptor results table.

### Data Export in AIRR Format

The improved standardization of receptor data fields also enables straightforward conversion of IEDB data into AIRR Rearrangement format (41). We have now implemented a direct export option in this format, supplementary to the existing customizable receptor data export, to facilitate interoperability with AIRR-compliant tools (52–56). The user can either query for the desired data subset and select ‘Export Type = AIRR’, or find the full data exports in AIRR format on the ‘Database Export’ page (iedb.org/database_export_v3.php).

The IEDB-AIRR export format aligns with the AIRR Rearrangement format with the addition of custom columns, recognizable by their ‘iedb_’ prefix. These custom columns describe epitope, MHC, and assay identifier data (see **Figure 6**). Each receptor-epitope pair is represented on separate rows, as well as separating positive and negative assay results. The AIRR Rearrangement format contains one row per chain, and uses the ‘cell_id’ field to pair together the receptor chains. Therefore, the IEDB-AIRR cell_id contains a conglomeration of receptor identifier, epitope identifier, and assay status (positive/negative), to ensure ‘cell_id’ occurs exactly once or twice in the IEDB-AIRR export, representing single-chain or paired-chain data, respectively. The AIRR field ‘iedb_receptor_group’ contains an identifier which can be traced back to its receptor details webpage by adding the ‘iedb.org/receptor/’ prefix. Note that multiple distinct receptors may belong to the same receptor group. Similarly, the identifier under ‘reactivity_ref’ can be combined with ‘iedb.org/epitope/’ to find the epitope details page.

**Figure 6:**
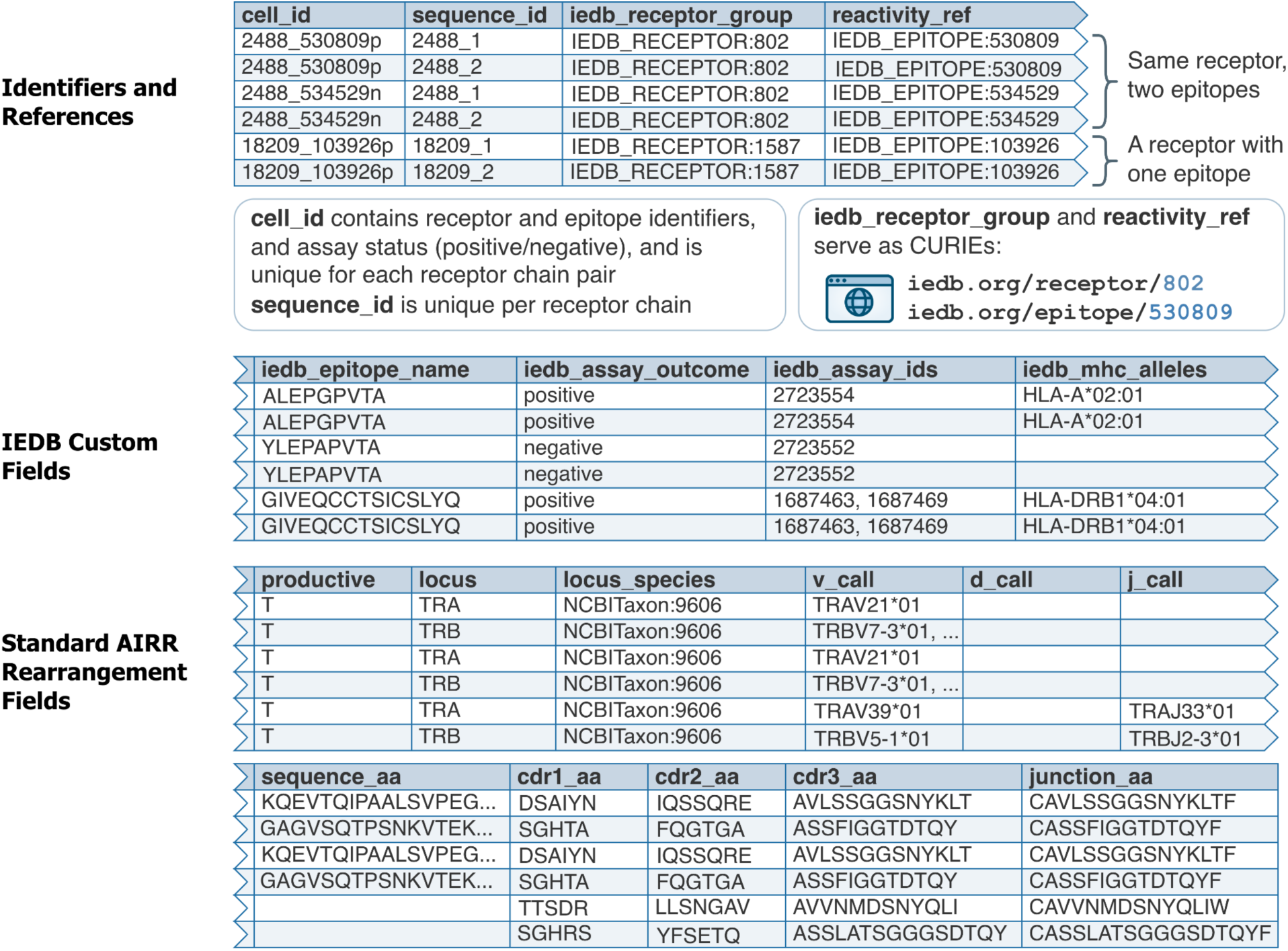
Example of AIRR export format in the IEDB. Each line describes a single receptor chain in interaction with one epitope. The cell_id can be used to pair together the two chains of a receptor specific to one epitope. The first four lines show the same receptor (receptor group 802) which has a positive assay for epitope ALEPGPVTA, and a negative assay for epitope YLEPAPVTA. The last two lines show a different receptor in combination with a single epitope, described in two different assays (1687463, 1687469). The epitope and receptor group IDs are Compact URIs (CURIEs) with associated summary webpages (e.g., iedb.org/receptor/802 or iedb.org/epitope/530809). Additional mandatory AIRR Rearrangement columns are omitted from this example but included in the actual export.

## Future Work

The IEDB aims to provide a comprehensive collection of epitopes and their experimentally validated cognate immune receptors, curated from the scientific literature. To address inconsistencies in receptor data formatting, nomenclature inconsistencies and missing values, we revised our computational processing pipelines and present a fully annotated, computationally validated and analysis-ready receptor dataset.

Although we have improved the accuracy and consistency of receptor sequence annotations, it should be noted that the IEDB does not assess or make judgments regarding the quality of receptor-epitope binding evidence, resulting in mixed-quality data. For instance, some 10X Genomics single-cell datasets are known to contain a large number of false positive binders (57, 58), and excluding these datasets can boost the performance of TCR specificity prediction models (12). While recent work has aimed to reduce noise in single-cell pMHC readouts through statistical analysis (59), certain data subsets remain noisy. Strikingly, a recent study experimentally validated the reactivity of hundreds of TCRs towards two epitopes in VDJdb, and were unable to confirm reactivity for half the tested TCR-pMHC pairs (60). The confidence scores assigned to these binding pairs by VDJdb (43) could not be used as a reliable indicator of true TCR reactivity (60). While generation of high-quality independently validated epitope binding datasets is highly valuable, it is costly, resource-intensive and has limited scalability compared to the full documented collection of receptor-epitope pairs. In future work, computational models may be used to identify and filter out potential low quality binding pairs in the IEDB.

Besides binding accuracy, receptor sequence annotation presents additional challenges. The IMGT reference sets have been widely adopted as the standard reference sets for immune receptor germline genes (61). However, these reference sets are known to be incomplete, biased towards Caucasian genotypes (62), may contain erroneous sequences (47, 63) and are subject to continuous updates, renaming and removal (64). Alternatively, through OGRDB the AIRR-C provides high-quality version-controlled IG reference sets (for human, mouse and macaque) containing only alleles supported by strong evidence (47), and a complementary human TR reference set is in preparation (30). We have transitioned to using the AIRR-C human IG reference sets, and plan to do so for any IG/TR reference set in the future. A persistent challenge, however, is that IG and TR gene naming is a live process, in which names and the sequences attached to those names can change in the light of new data (33). This underscores the importance of the reference set versioning and transparency enabled by OGRDB as per FAIR principles (30, 65), and the IEDB will update the gene names it uses as the community consensus on nomenclature evolves.

## Data availability statement

All curated and calculated receptor datasets are available on iedb.org. Weekly builds of the full datasets can be found at https://www.iedb.org/database_export_v3.php (tcr_full_v3.xlsx, bcr_full_v3.xlsx and airr_full.zip). All results presented in this manuscript are generated based on the 05/28/2026 data export.

## Conflict of interest statement

B.P. is a consultant for Mint Precision Analytical, Amgen Inc. and Sanofi, and a Scientific Founder of Lykeion. B.C. is a partner in iReceptor Genomic Services. L. G. C. is Chief Scientific Officer for OncoSeer Diagnostics.

## Funding

The IEDB is funded by the National Institute of Allergy and Infectious Diseases [75N93019C00001, 75N93026C00001]; CEDAR is funded by the National Cancer Institute [U24CA248138]; the AKC is funded by the National Institute of Allergy and Infectious Diseases [U24I177622].

## Notes

https://www.iedb.org/database_export_v3.php

