## Supplementary Tables and Figures for "Revised Adaptive Immune Receptor Data in the Immune Epitope Database"

**Supplementary Table 1: The number TCRs per species in the IEDB**

| Species | Number of distinct TCRs | Number of TCR groups |
|---|---|---|
| Homo sapiens | 200,555 | 177,664 |
| Mus musculus | 7,325 | 6,992 |
| Synthetic construct | 2 | 2 |
| Macaca mulatta | 1 | 1 |
| Gallus gallus | 1 | 1 |

**Supplementary Table 2: The number BCRs per species in the IEDB**

| Species | Number of distinct BCRs | Number of BCR groups |
|---|---|---|
| Homo sapiens | 3,087 | 2,801 |
| Mus musculus | 1,320 | 1,161 |
| Lama glama | 295 | 285 |
| Vicugna pacos | 169 | 165 |
| Camelus dromedarius | 97 | 83 |
| Oryctolagus cuniculus | 65 | 61 |
| Macaca mulatta | 27 | 27 |
| Rattus norvegicus | 26 | 24 |
| Camelidae | 25 | 23 |
| Gallus gallus | 23 | 16 |
| Macaca fascicularis | 18 | 18 |
| Camelus bactrianus | 13 | 11 |
| Bos taurus | 9 | 9 |
| Sus scrofa | 5 | 5 |
| Nothocricetulus migratorius | 4 | 4 |
| Synthetic construct | 4 | 4 |
| Squalus acanthias | 3 | 3 |
| Ginglymostoma cirratum | 3 | 2 |
| Orectolobus maculatus | 3 | 2 |
| Lama | 2 | 2 |
| Rattus | 2 | 2 |
| Chiloscyllium plagiosum | 2 | 2 |
| Camelus ferus | 2 | 2 |
| Lepus | 1 | 1 |
| Mus | 1 | 1 |
| Myodes glareolus | 1 | 1 |
| Chlorocebus aethiops | 1 | 1 |
| Pan troglodytes | 1 | 1 |

**Supplementary Table 3: The number of curated TCR chains in the IEDB with either full nucleotide/amino acid sequence information or CDRs with gene names.**

| Curated Data type | TCR α | TCR β | TCR γ | TCR δ | BCR heavy | BCR light |
|---|---|---|---|---|---|---|
| CDRs and gene names (no full sequence) | 57,047 | 178,662 | 236 | 155 | 1,449 | 597 |
| Full VDJ sequence - amino acid | 588 | 613 | 5 | 5 | 3,745 | 3,006 |
| Full VDJ sequence - nucleotide | 136 | 116 | 0 | 0 | 81 | 66 |
| Total across data types | 57,771 | 179,391 | 241 | 160 | 5,275 | 3,669 |

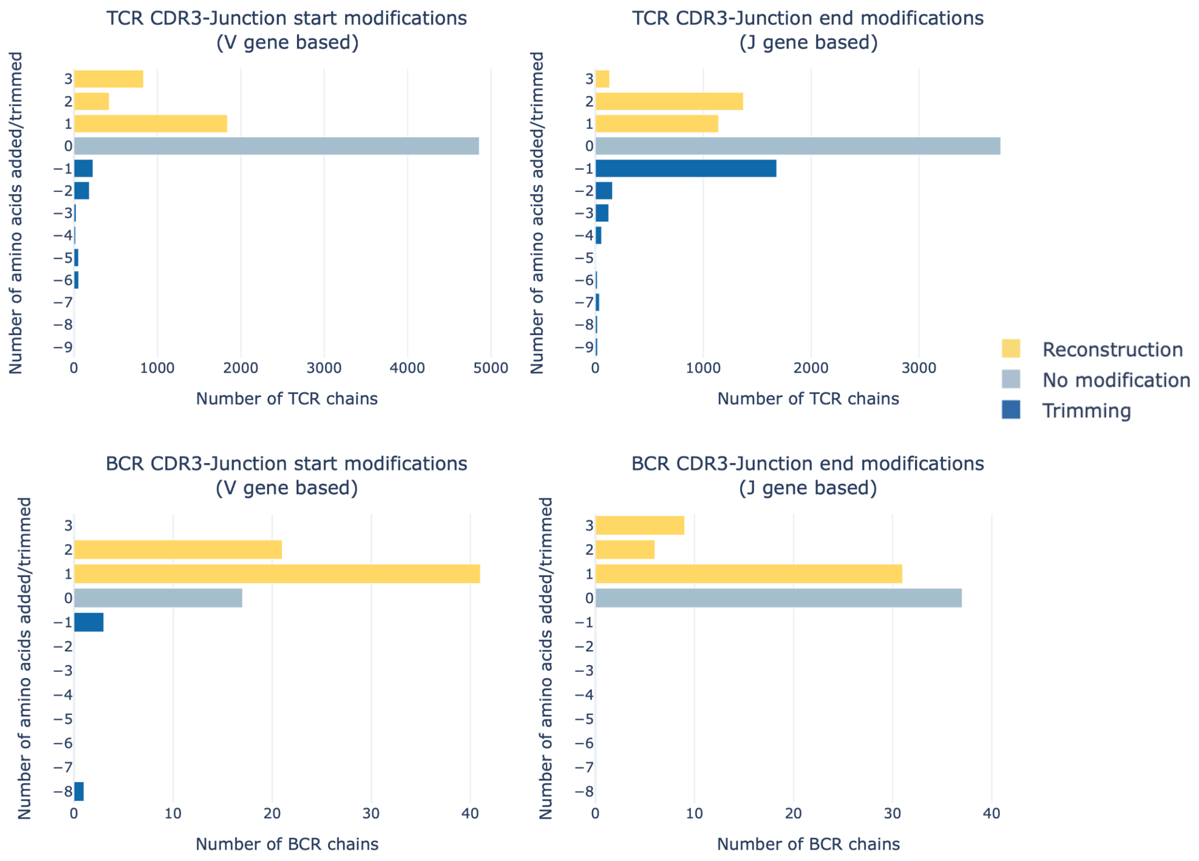


**Supplementary Figure 1: The number of amino acids reconstructed or trimmed from the curated CDR3 to correct the CDR3 Junction sequence.** The maximum number of reconstructed residues was capped at 3 when V/J gene information was available, and 1 (only constructing the conserved residue) when no V/J gene was supplied. Note that the CDR3 sequences which already corresponded exactly to the IMGT Junction or IMGT CDR3 definition (including or excluding the conserved residues) are excluded here (see instead **Figure 3a**). Thus, no modification at the start of the CDR3 means there was some modification at the end, and vice versa. The most common modifications are one addition on the V side (missing Cys104) or removal of one extra amino acid on the J side (ending with 'FG'). Corrections were computed with tidytcells v3.0.0-alpha (github.com/yutanagano/tidytcells/tree/dev/v3) (39).

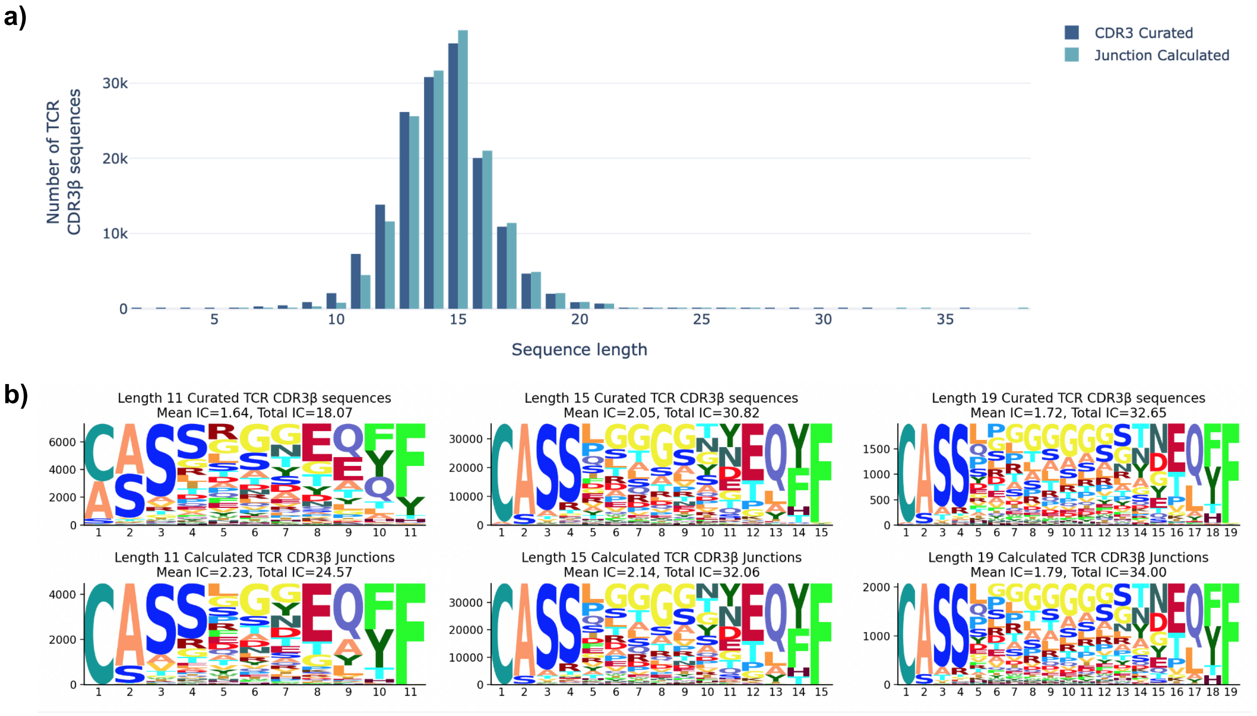


**Supplementary Figure 2: Comparison of the TCR CDR3β sequences before and after standardization.** a) A histogram of all unique (deduplicated) sequences before and after standardization. The *calculated* CDR3 Junction sequences are slightly longer due to a substantial fraction of the *curated* CDR3 values not containing the conserved flanking amino acids. b) Logo plots are shown for three different sequence length groups from the histogram given in a. The sequence diversity in the conserved flanking regions is reduced after standardization (bottom row), due to the conserved anchoring residues being aligned. This effect is strongest in the shorter CDR3 sequences.

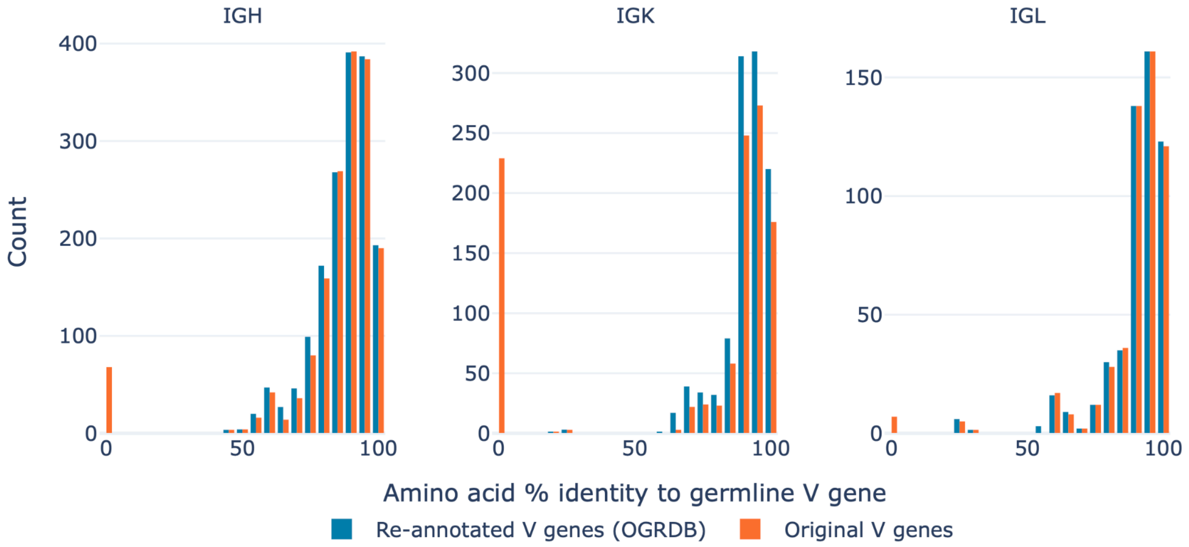


**Supplementary Figure 3: Mutations to germline V gene in human antibody data is significantly lower in re-annotated data.** Sequence identity was calculated in the germline-encoded (up to Cys104) positions of the full variable sequences of human antibodies, and compared to the germline sequence of the assigned V allele. The original annotations were compared to the IMGT human IG reference set, whereas the re-annotated data was compared to the same AIRR-C human IG reference set used for V gene assignment, and 0% sequence identity was assigned when a gene did not exist in the given reference set. The updated annotations produce a significantly higher sequence identity to germline, with average IGHV, IGKV and IGLV identity values of 87.1% vs. 82.9%, 91.1% vs. 70.4% and 90.2% vs. 88.5% for the updated annotations vs. previous, respectively. In particular for kappa light chains, 23.3% of V gene annotations in the original data did not exist in the human IG reference set.

a)

b)

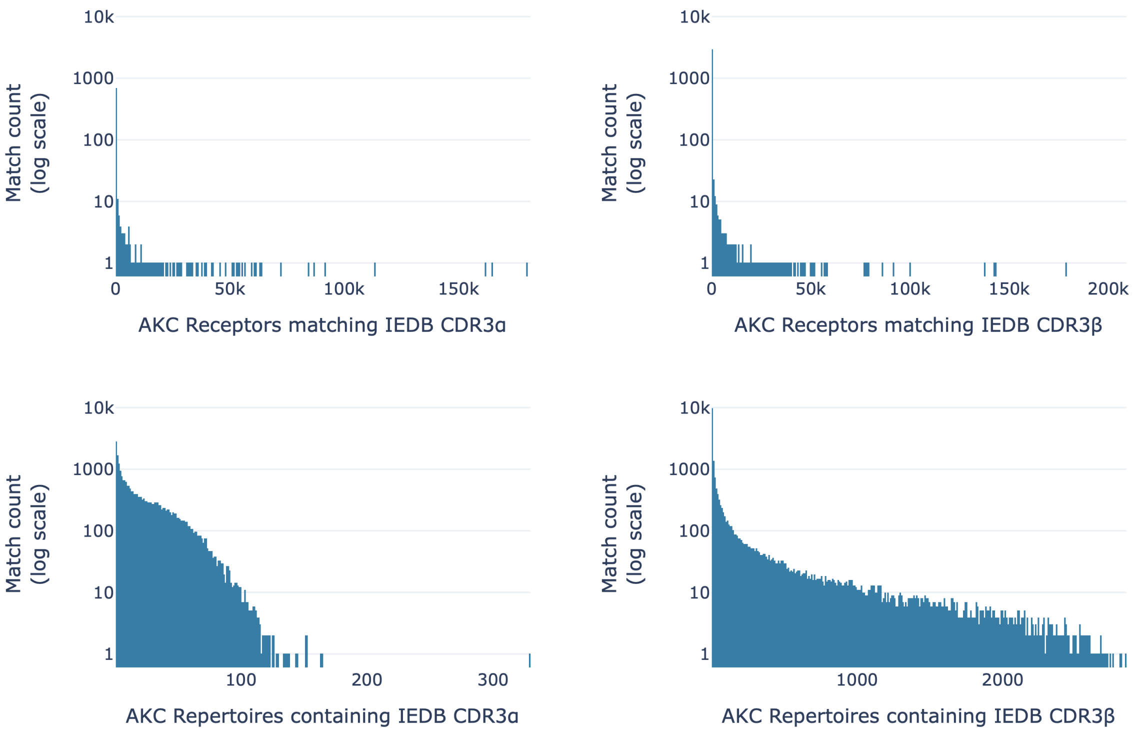


**Supplementary Figure 4: Number of α and β chain CDR3 matches to the AKC.** a) The distribution of match counts shows few IEDB receptor groups having CDR3 matches to up to as many as 150,000 ~ 200,000 unique receptors in the AKC, and hundreds to thousands of IEDB receptor groups with at least one respective CDR3 α or β chain match in the AKC. b) The IEDB receptor groups can be found across hundreds of different α chain repertoires and thousands of β chain repertoires in the AKC.

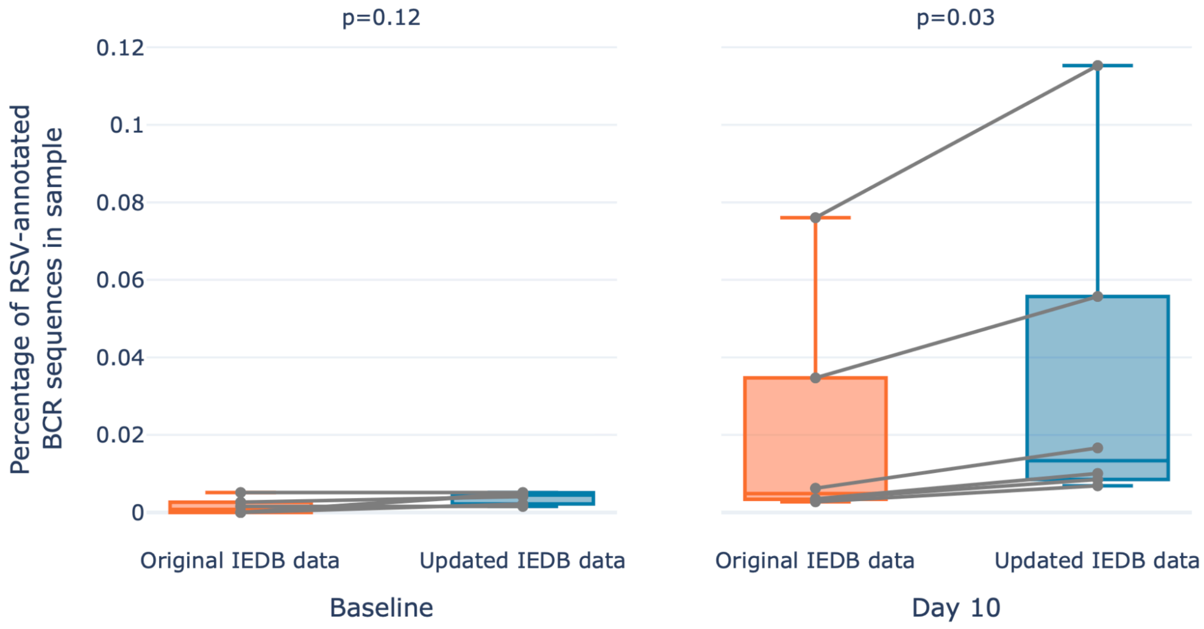


**Supplementary Figure 5: Increased sensitivity due to increased completeness of updated database.** Sequences from six subjects at baseline and 10 days post-RSV challenge were obtained from (51), and specificity prediction was performed using CloneSearch (23) against RSV-specific human antibodies from the IEDB (70% CDRH3 identity cut-off and use_alleles set to False). For comparison with the updated IEDB data, IGHV genes were re-annotated using the same AIRR-C IGHV reference set. The updated IEDB data yields significantly more RSV-specific predictions at day 10 compared to baseline (Wilcoxon signed-rank test).
